# The Rosenbluth sampling Calculation of Hydrophobic-Polar Model

**DOI:** 10.1101/2022.05.02.490260

**Authors:** Marcin Wierzbiński, Alessandro Crimi

## Abstract

Lattice proteins are models resembling real proteins. They comprise an energy function and a set of conditions specifying the interaction between elements occupying adjacent lattice sites. In this paper we present an approach examining the behavior of chains of a large number of molecules. We investigate this by solving a restricted random walk problem on a cubic lattice and square lattice. More specifically, we apply the *hydrophobic-polar* model to examine the spatial characteristics of protein folds using the Monte Carlo method. This technique is the so-called Rosenbluth sampling method for solving restricted random walk problems. Specifically, by solving such walks we obtain plausible folds. In addition, this method can be extended to solve the hydrophobic-polar model. In this paper, we describe this method as an algorithm that calculates the energy spectrum for the hydrophobic-polar model, and the related formula for estimating the number of folds. Moreover, we estimate the number of folds for each sequence using hydrophobic-polar model energy estimation. On test sequences the predicted protein folds were obtained with a mismatch of one unit according to the energy. We also observe that the estimated number of folds depends only on the length and not on the type of sequence. This promising strategy can be extended to quantify other proteins in nature.

## 1 Introduction

The search for a more efficient algorithm of protein folding in the hydrophobic-polar (HP) model is an important aspiration in many disciplines (Sali et al. (1994), Pande (2010)). Knowing how proteins fold can help elucidate their three-dimensional structure-function relationship, which is crucial to the understanding of enzymes and to the treatment of misfolded-protein diseases such as Alzheimer’s, Huntington’s, and Parkinson’s disease. The numerical simulation focused on those proteins is particularly useful for drug design, as it allows to test different physical characteristics using models of various complexities. Indeed, if high-resolution chemical structure is used, leading to precise molecule representations, dynamical simulation showing atomic interactions can be reached. This might ultimately provide more effective and personalized drugs.

It has been shown that the HP protein folding model is NP-Hard (Berger & Leighton (1998)), which means it is difficult to solve efficiently for longer protein sequences. In order to overcome this obstacle, many heuristic algorithms have been proposed (Jiang et al. (2003), Yanev et al. (2017)). Besides heuristics mostly based on optimization, other approaches are based on the idea that cooperativity of folding occurs, as local conformational choices which constraints the optimization space in which solutions are searched. Those assumption-based methods include hydrophobic zipper method Dill et al. (1993), which assumes that once a hydrophobic contact is created it cannot be broken. And the core-directed chain growth method Beutler & Dill (1996) which constrains the optmization search within the space of solutions having a hydrophobic core with a square (in 2D) or a cube (in 3D).

In this context, there is theoretical and experimental evidence of the advantage of solving a restricted random walk problem (RRW) on cubic and square lattices. One of the earliest proposed numerical algorithms which apply the RRW paradigm is the one designed by M. Rosenbluth and A. Rosenbluth (Rosenbluth & Rosenbluth (1955)). In this report, we present a benchmark implementation of Rosenbluth methods for the HP model with an additional extension to estimate the number of possible sequence configurations.

### 1.1 Hydrophobic-Polar model

In the hydrophobic-polar model, the set of twenty standard amino acids is reduced to two: *H* (hydrophobic amino acid) and *P* (hydrophilic amino acid). More formally, the model relies on *embedding* a given finite polypeptide *sequence s* = (*s*_1_, …, *s*_*i*_, …, *s*_*k*_) where *s*_*i*_ ∈ {*H, S*}, into a given infinite graph *G*. In this article, the graph *G* will primarily be the three-dimensional cubic lattice *G* = ℤ^3^ and square cubic lattice *G* = ℤ^2^ over integer numbers ℤ. A *fold* of length *k* for *s* in *G* is an injective mapping *f* : [1, …, *k*] ⟼ *G* such that adjacent integers map to adjacent points of *G*. The set of all folds of length *k* is denoted as 𝒵_*k*_. In addition, each point *f* (*i*) is assigned one letter from the polypeptide sequence *s*_*i*_. Such neighboring points form a *bond*. Each point of ℤ^3^ has six neighbors (*x* ± 1, *y* ± 1, *z* ± 1). The energy of the fold of *s* is expressed as

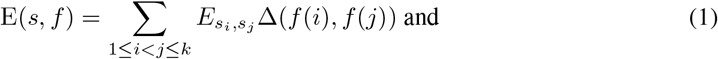

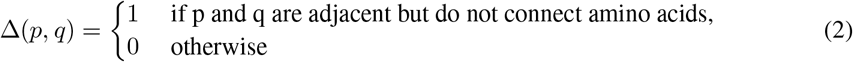

with energy equation:

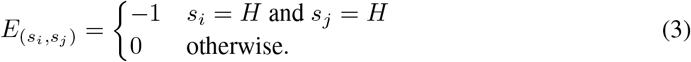

The above equation for calculating the energy of fold *s* in *G* can also be expressed as negation of the number of *H* − *H* bonds in the fold, where a bond is a pair of symbols corresponding to adjacent points, except for those *H*’s which are adjacent to pairs of sequences *s*. The goal of the HP model is to minimize the energy

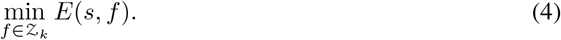

### 1.2 Rosenbluth sampling method

Note that for a given number of adjacent points *k* in the fold, any configuration consisting of *k* adjacent points laid out joined in succession on a cubic lattice ℤ^3^ is considered. The method proposed by Rosenbluth & Rosenbluth (1955) involves drawing successive steps of a random walk only from among *acceptable points*, which are points previously not visited. In this section, we describe the random procedure in more detail. We will focus on the 3D case, i.e *G* = ℤ^3^, but the method is easily transferable to the 2D case.

Regarded as a random walk problem, for any walk consisting of *m* adjacent points and ending at position (*x, y, z*)_*m*_, all six positions are a priori equally likely at iteration *m*. The excluded volume effect is simulated by the requirement that a fold is not allowed to cross itself or back up on itself at any iteration. Consequently, at any iteration, there are at most five possible positions to move to. For simplicity, we assume that the first link originates from (0, 0, 0). Any satisfactory set of *m* adjacent points start from the origin (*x, y, z*)_*m*_ is associated with a weighting function *w*_*m*_ of possible positions calculated at each step according to the procedure described below. At any iteration *m* where the most recent link terminates at (*x, y, z*)_*m*_ and 5 potential positions (*x*±1, *y*±1, *z*±1)_*m*_ must be considered, while position (*x, y, z*)_*m*−1_ is ruled out immediately. All five remaining potential positions at *m* + 1 may be associated with values (*x, y, z*)_*i*_ for *i* = *m* − *p* where *p* is an odd number greater than 1. If the comparison reveals this to be the case, a modification of the weight *W*_*m*_ must be made, obtaining *W*_*m*+1_.

Below we present all possible cases at iteration *m*:

1. All six new position (*x*±1, *y*±1, *z*±1)_*m*_ are occupied. The process is then terminated with weight *W*_*m*_ = 0.
2. Only *w*_*m*_ new positions are unoccupied, with 0 < *w*_*m*_ ≤ 5. Then

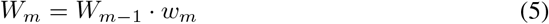

During this process, an embedding is generated. If the embedding is equal to the length of the sequence, we can calculate energy according to the presented formula.

#### Estimation of fold numbers

In this section, we present a mathematical justification for the estimation of folds.

Let us assume in general terms that when constructing fold *f*_*i*_ of length *k* with partial weights 0 < *w*_*i*_ ≤ 5:

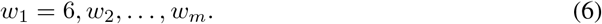

It is not excluded that at a certain step we may have no further possibilities for continuation, i.e., *w*_*m*+1_ = 0. We then say that a *non-extendable* fold of length *m* has been formed.

Let 𝒴 denote the set of all folds of length *m* = *k* and non-extendable folds of length *m* < *k*. Recall that set of all folds of length *m* = *k* is denoted as 𝒵_*k*_. Clearly, 𝒵_*k*_ *⊂* 𝒴.

The probability of picking a random fold *f*_*i*_ ∈ 𝒴 of length *m* is equal to:

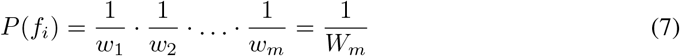

with a weight function for the specific fold *f*_*i*_. Let *f*_*i*_ ∈ 𝒴, so

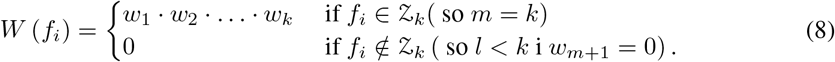

One can interpret *W* (*f*_*i*_) as the weight of fold *f*_*i*_. Let us now repeat the draw using the growth method *n* times. There are *n* random folds *f*_1_, *f*_2_, …, *f*_*n*_ from set *Y*.

Let *n*_*s*_ denote the number of drawn elements *f*_*i*_ for which *W* (*f*_*i*_) = *s* and the set of these elements 𝒲_*s*_ = {*f* ∈ 𝒴 : *W* (*f*) = *s*}. Then, based on the large numbers law:

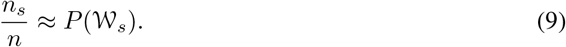

Therefore, the above expression can be written as the average weight of the drawn folds. We note that:

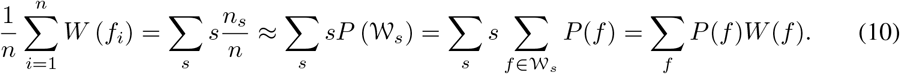

Finally, the expression for 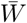 can be written as:

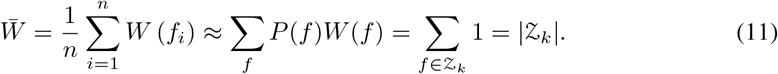

We introduce the following notation for fold estimators of length k:

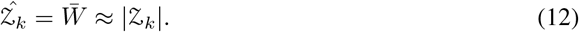

To validate this fold estimator we test sequences of different length and type and the results are reported in the following section.

#### Estimation standard error for 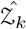

For estimation of standard error for 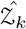, we used the *batch means method*. We will briefly describe how this method was applied in this setup.

Let us divide the sequence of weights:

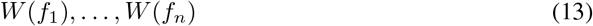

of length *n* into *j* ”blocks” of length *l* each (so n = *jl*):

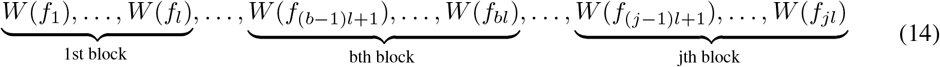

Let us denote by 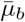 the mean calculated from the block

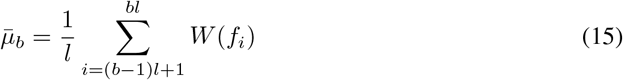

The estimator for the variance is defined as:

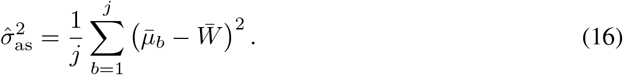

Then the standard error is the root of the estimator for the variance:

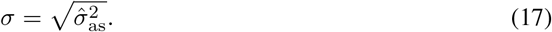

This method significantly speeds up the variance calculation by the standard method of generating estimators and calculating the standard deviation.

## 2 Results

The experiments were run on Google’s Colab platform on Intel(R) Xeon(R) CPU @ 2.20GHz with 13GB RAM. We investigated 2 different datasets one 2D and one 3D. The algorithm code, written in Python, can be found at the following website: Wie (2022). The method was run for *n* = 10^5^ of suitable configurations folds with a specific sequence *s* of length *k*. We used *j* = 10^3^ blocks in the batch means method for estimation standard error.

### 2.1 Benchmark for 2D

Using the method proposed above, we calculate 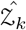 with a statistical error. The algorithm was initially tested for several sequences in dimension 2 (for ℤ^2^ from site LABORATORY (2011).

This first experiment lasted 5 minutes. Symbols *H*^*i*^ and *P*^*i*^ in the table correspond to *i* repetitions of sequence characters. The results related to this dataset are reported in Table 1, with 4 examples of resulting predicted folding depicted in Figure 1.

**Table 1:**
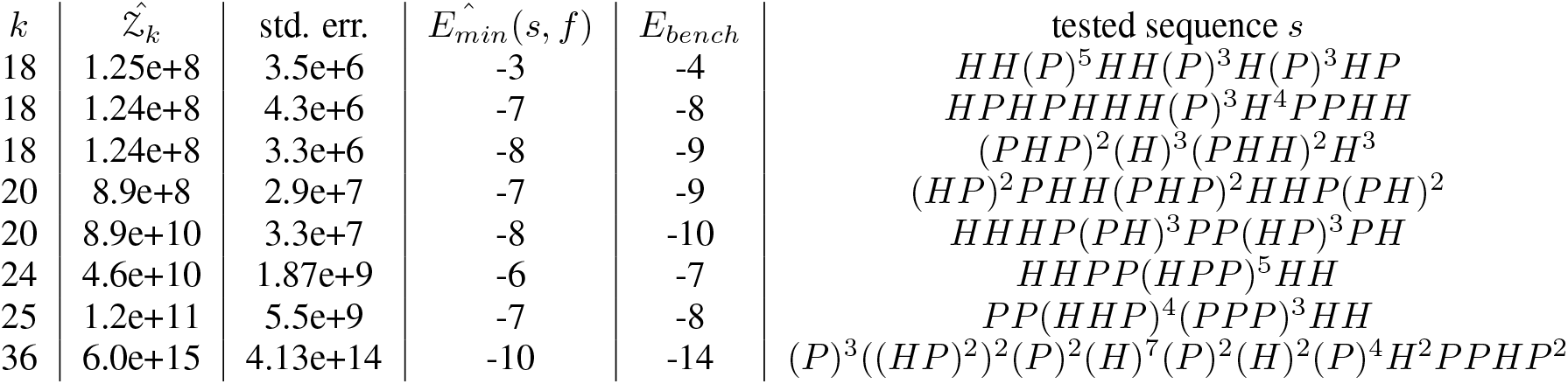
In this table we compare how our method performs against the model from LABORATORY (2011). We can conclude that our energy is minimally different. Having computed an estimation for all folds ℤ^2^ for each sequence, we can conclude that the number of folds 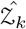 does not depend on the tested sequence *s*. We can observe that the estimated number of folds depends only on length *k* and not on the form of *s*.

**Figure 1:**
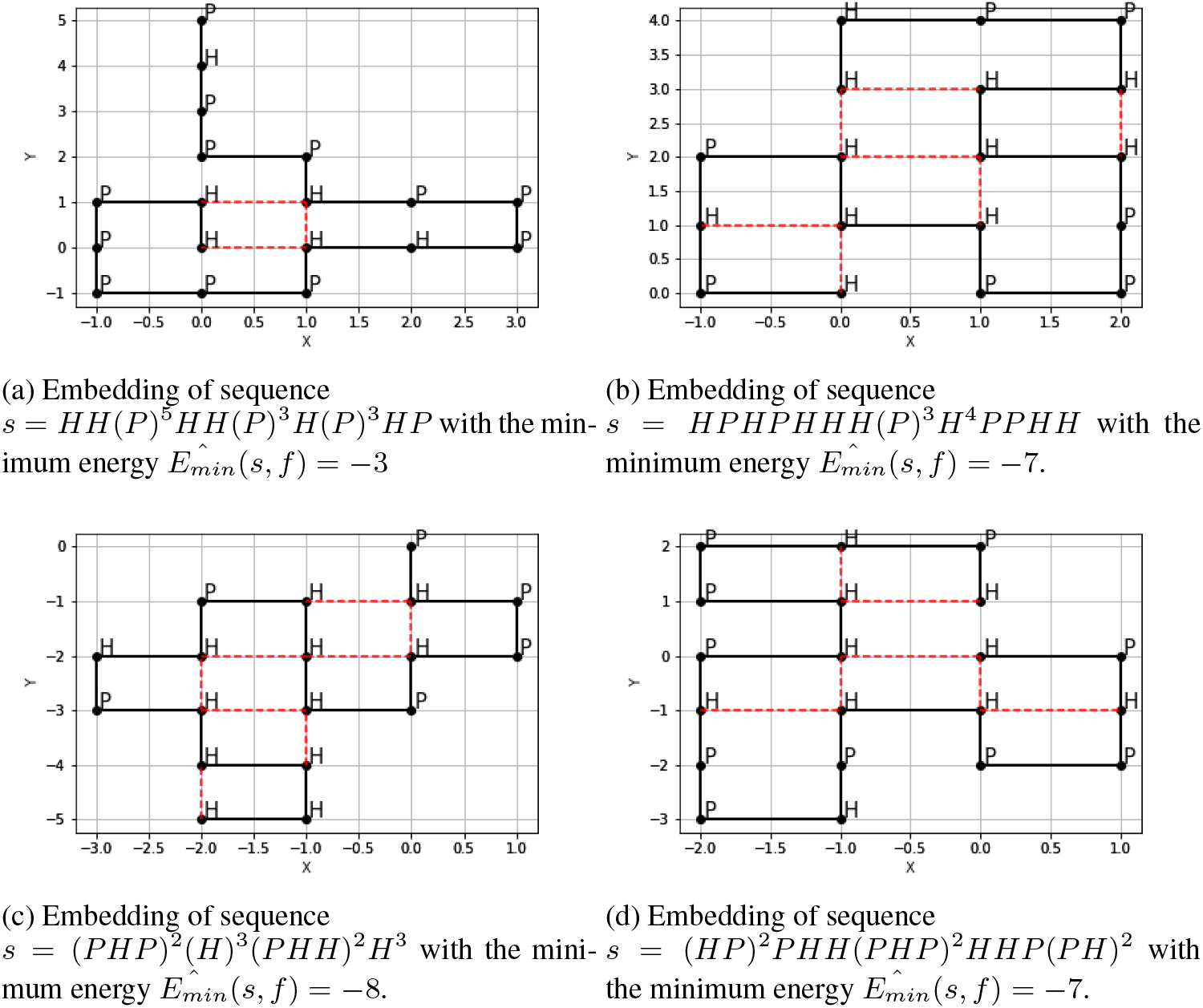
In each figure we embed a given finite polypeptide sequence in the square lattice *G* = ℤ^2^. The energy in the presented diagrams can be easily deduced. Each red line indicates a bond. The number of these edges corresponds to energy 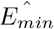.

### 2.2 Estimation for 3D

The experiments were performed using *n* = 10^5^ and *j* = 10^3^ blocks in batch means method. The code can be found in Wie (2022). Estimated energy is equal to 0 for all *k*. This is because it is difficult to fold for ℤ^3^ so that the number of *H* − *H* bonds is minimised. The second experiment lasted 4 minutes. Results for this dataset are reported on Table 2, with 2 examples of resulting predicted folding depicted in Figure 2.

**Table 2:** In the accompanying table we count the fold estimation values for dimension 3 for ℤ^3^. Referring to experiment 1 we see that there is a significant increase in the number 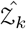 of these folds for each sequence. For values of *k* = 24, 25 we observe particularly large differences.

| $k$ | estimation $\hat{\mathcal{Z}}_k$ | std. err. | tested sequence $s$ |
| --- | --- | --- | --- |
| 1 | 6 | 0 | $H$ |
| 2 | 30 | 0 | $HP$ |
| 3 | 150 | 0 | $HPH$ |
| 4 | 726.0 | 1.6 | $HPHP$ |
| 5 | 3532.1 | 11.14 | $HPPHP$ |
| 6 | 16934.8 | 69.62 | $HPPHPH$ |
| 10 | 8.8e+6 | 6.88e+4 | $HH(P)^4(H)^3P$ |
| 18 | 2.23e+12 | 3.5e+10 | $(H)^2(P)^5(H)^2(P)^3H(P)^3HP$ |
| 20 | 4.96e+13 | 8.3e+11 | $(HP)^2PH(HP)^2(PH)^2HP(PH)^2$ |
| 24 | 2.47e+16 | 5.2e+14 | $P(PH)^3(PP)^2(HH(PP)^2)^2H$ |
| 25 | 1.16e+17 | 2.25e+15 | $P(PH)^3(PP)^2(HH(PP)^2)^2HH$ |

**Figure 2:**
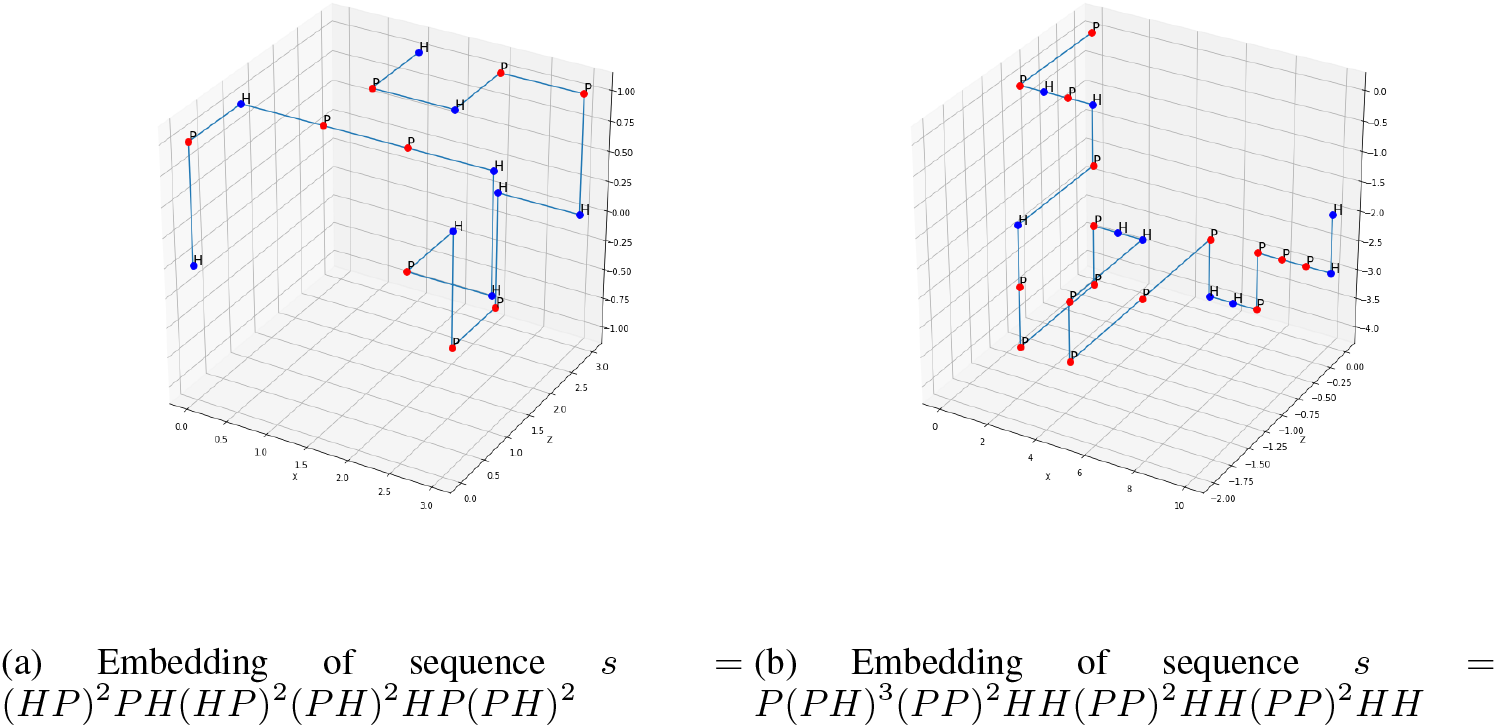
In the graph above, we can observe that the fold did not wrap in such a way that the two *H* − *H*s are next to each other. Therefore, energy is equal to 0. This is because we have significantly more degrees of freedom in 3D space.

The experiment itself shows how difficult it is to wrap sequences in 3 dimensions. Estimations for sequences 24 and 25 alone show that the number of folds is on the order of 10^16^ and 10^17^, as shown in Table 1 and 2. However, the energy of the fold is still zero. Therefore, the 3D model is significantly more difficult than the 2D model.

For the initial cases *k* = 1, 2, the results are obvious. If the length of the sequence is 1 (*k* = 1), then we have only 6 possible points. For *k* = 2 we have 36 possible point assignments, 6 of which are forbidden. We have also prepared special sequences for cases *k* ∈ {1, 2, 3, 4} that will always have zero energy. It is not possible to wrap a sequence in such a way that points with *H* − *H* labels touch each other.

## 3 Discussion

Correctly predicting protein conformations based on the amino acid sequence is of pivotal importance for drug design and other relevant computational chemistry tasks. In this paper, we report our computational experiments, where we use HP sequences corresponding to published benchmarks LABORATORY (2011) with a 2D lattice in the HP model. Our model successfully estimates the number of folds for a particular sequence; regardless of the type of sequence but only on its length. For small sequences, the method accurately estimates the number of folds. Our experiments show that for sequences of size *k* = 24, 25 the 3D model becomes significantly more complex than the 2D model. It has been observed that adding one dimension significantly affects the solution base. In 2D, the energies are −6 and −7, respectively, while in 3D the energy is zero for both cases. There are too many degrees of freedom to draw consecutive points. Therefore, it is difficult to find a wrap that has non-zero energy even for shorter sequences. However, the Rosenbluth sampling method can be successfully used to estimate the number of all folds, especially those with energy 0. This can help design heuristic algorithms based on this hindsight. The estimation itself, according to our mathematical justification, increases in accuracy as we increase the number of iterative executions of the method. The described approach is effective for identifying and sampling configurations on a lattice geometry. This kind of representation can be useful in the context of ab initio protein structure prediction Rashid et al. (2016). Expansions as implementations on quantum devices have been proposed, but those have been limited to the 2D case so far Micheletti et al. (2021). Conversion of the proposed tool into quadratic unconstrained binary optimization (QUBO) Kochenberger et al. (2014) using 3D lattices on quantum devices will be investigated in future work.

## 3.1 Acknowledgements

This research is supported by the European Union’s Horizon 2020 research and innovation programme under grant agreement Sano no 857533, and by the International Research Agendas programme of the Foundation for Polish Science, co-financed by the European Union under the European Regional Development Fund.

## References

Monte Carlo Calculation of Protein Folding. https://github.com/marcin119a/Monte-Carlo-Calculation-of-Protein-Folding, 2022. [Online; accessed 02-Feb-2022].

B Berger and T Leighton. Protein folding in the hydrophobic-hydrophilic (hp) model is np-complete. Journal of computational biology, 5(1):27–40, 1998. ISSN 1066-5277.

Thomas C Beutler and Ken A Dill. A fast conformational search strategy for finding low energy structures of model proteins. Protein Science, 5(10):2037–2043, 1996.

Ken A Dill, Klaus M Fiebig, and Hue Sun Chan. Cooperativity in protein-folding kinetics. Proceedings of the National Academy of Sciences, 90(5):1942–1946, 1993.

Tianzi Jiang, Qinghua Cui, Guihua Shi, and Songde Ma. Protein folding simulations of the hydrophobic–hydrophilic model by combining tabu search with genetic algorithms. The Journal of chemical physics, 119(8):4592–4596, 2003.

Gary Kochenberger, Jin-Kao Hao, Fred Glover, Mark Lewis, Zhipeng Lü, Haibo Wang, and Yang Wang. The unconstrained binary quadratic programming problem: a survey. Journal of combinatorial optimization, 28(1):58–81, 2014.

ISTRAIL LABORATORY. HP 2D Benchmarks. https://www.brown.edu/Research/Istrail_Lab/hp2dbenchmarks.html, 2011. [Online; accessed 02-Feb-2022].

Cristian Micheletti, Philipp Hauke, and Pietro Faccioli. Polymer physics by quantum computing. Physical Review Letters, 127(8):080501, 2021.

Vijay S Pande. Simple theory of protein folding kinetics. Physical review letters, 105(19):198101–198101, 2010. ISSN 0031-9007.

Mahmood A Rashid, Sumaiya Iqbal, Firas Khatib, Md Tamjidul Hoque, and Abdul Sattar. Guided macro-mutation in a graded energy based genetic algorithm for protein structure prediction. Computational biology and chemistry, 61:162–177, 2016.

Marshall N Rosenbluth and Arianna W Rosenbluth. Monte carlo calculation of the average extension of molecular chains. The Journal of chemical physics, 23(2):356–359, 1955. ISSN 0021-9606.

Andrej Sali, Eugene Shakhnovich, and Martin Karplus. How does a protein fold? Nature (London), 369(6477):248–251, 1994.

Nicola Yanev, Metodi Traykov, Peter Milanov, and Borislav Yurukov. Protein folding prediction in a cubic lattice in hydrophobic-polar model. Journal of Computational Biology, 24(5):412–421, 2017.

